# Metagenomic profiling of host-associated bacteria from 8 datasets of the red alga *Porphyra purpurea*, with MetaPhlAn 3.0

**DOI:** 10.1101/2020.11.17.386862

**Authors:** Orestis Nousias, Federica Montesanto

## Abstract

Microbial communities play a fundamental role in the association with marine algae, in fact they are recognized to be actively involved in growth and morphogenesis.

*Porphyra purpurea* is a red algae commonly found in the intertidal zone with an high economical value, indeed several species belonging to the genus *Porphyra* are intensely cultivated in the Eastern Asian countries. Moreover, *P. purpurea* is widely used as model species in different fields, mainly due to its peculiar life cycle. Despite of that, little is known about the microbial community associated to this species. Here we report the microbial-associated diversity of *P. purpurea* in four different localities (Ireland, Italy United Kingdom and USA) through the analysis of eight metagenomic datasets obtained from the publicly available metagenomic nucleotide database (https://www.ebi.ac.uk/ena/). The metagenomic datasets were quality controlled with FastQC version 0.11.8, pre-processed with Trimmomatic version 0.39 and analysed with Methaplan 3.0, with a reference database containing clade specific marker genes from ~ 99.500 bacterial genomes, following the pan-genome approach, in order to identify the putative bacterial taxonomies and their relative abundances. Furthermore, we compared the results to the 16S rRNA metagenomic analysis pipeline of MGnify database to evaluate the effectiveness of the two methods. Out of the 43 bacterial species identified with MetaPhlAn 3.0 only 5 were common with the MGnify results and from the 21 genera, only 9 were common. This approach highlighted the different taxonomical resolution of a 16S rRNA OTU-based method in contrast to the pan-genome approach deployed by MetaPhlAn 3.0.

## 1. Introduction

Microbial communities are the keystone in all terrestrial and marine habitats, being involved in biogeochemical cycles, primary productivity, symbiotic relationships, metabolic pathways and so on (Konopka, 2009). In the last decades, metagenomic studies in the marine environment greatly enhanced the knowledge about the diversity and complexity of marine microbial communities allowing straight sequencing of DNA from environmental samples or directly from the hosts these communities are associated with (Biller et al., 2018). However, the biodiversity of microbial communities as well as their relationship with the environment, in terms of chemistry and interactions with its inhabitants is still scantly known (Biller et al., 2018).

The genus *Porphyra* C.Agardh, 1824 (Bangiales, Bangiophyceae) includes more than one hundred species of red algae worldwide distributed, with a remarkable economical value since they are cultivated in China, Korea and Japan as human food resource due to their contents of proteins, vitamins and minerals, worthing around $1.4 billion/year (Yarish and Pereira, 2008). Moreover, *Porphyra* is considered a model for basic and applied research, due to its unique dimorphic cycle, which consists of a gametophyte forming a large and leafy thallus, alternate with a filamentous sporophyte (Saga et al., 2002; Gantt et al., 2010). Thus, with regards to its unique life cycle, economical value, as well as for the significant resilience to high light and desiccation, *Porphyra* is considered also a model for genome sequencing (Gantt et al., 2010). Marine bacteria are actively involved in the normal growth and development of macro-algae, but only few studies have been carried on this regards.

Despite of the economical and scientific importance of *Porphyra* and the recognized implication of its microbial communities associated on its life cycle and not only, further studies are needed to deeply understand the composition and the relationship of the species belonging to the genus *Porphyra* with their microbiome.

Studies on *Porphyra yezoensis* have demonstrated that both the life cycle and growth of the species are markedly affected by associated or symbiotic bacteria. Indeed, in axenic cultures, *P. yezoensis* loose its typical morphogenesis during the gametophytic phase, although it kept morphogenesis during the sporophytic phase; but adding specific symbiotic bacteria to the xenic cultures, *P. yezoensis* recovered the deficient morphogenesis, restoring the typical folious morphology of the gametophytic generation. These results lead to the conclusion that these symbiontic bacteria are needed for the typical growth and morphogenesis of the thallus (Mori et al., 2004; Tang et al., 2014). In particular Namba et al., (2010) analysed the 16S rDNA gene from cultured strains of *P. yezoensis* finding sequences of several bacteria characterized into six groups comprising α-, *β*-, and *γ-proteobacteria, Lentisphaerae, Sphingobacteria*, and *Flavobacteria*.

On the other hand, since the reproductive phases among the *Porphyra s*pecies can vary greatly, it is likely that this could be reflected on the composition of the microbial species associated to each different species belonging to this genus (Brodie and Irvine, 2003). It is also important to underline that different species of macroalgae collected from the same area support different bacterial communities, while the same species are often characterized by similar bacterial communities even if collected from different localities, which can vary among each species at season level (Lachnit et al., 2011).

To date, the most widely for characterizing the diversity of microbiota is amplicon sequencing. This strategy is based on targeting and amplifying by PCR a taxonomically informative genomic marker that is common to all organisms of interest. The resultant amplicons are sequenced and characterized to determine which microbes are present in the sample and at what relative abundance. In the case of Bacteria and Archaea, amplicon sequencing studies target the small and large subunit of ribosomal RNA (16S) locus, which is an informative marker both for phylogeny and taxonomy analyses (Pace et al., 1986; Hugenholtz and Pace, 1996). Amplicon sequencing of the 16S locus has given insights on microbial diversity on Earth (Pace, 1997; Rappé and Giovannoni, 2003; Lozupone and Knight, 2007) and used to characterize the biodiversity of microbes from a range of environments including ocean thermal vents (McCliment et al., 2006), hot springs (Bowen De León et al., 2013), antarctic volcano mineral soils (Soo et al., 2009).

Moreover, amplicon sequencing is not without limitation as it may not achieve a resolution of a substantial fraction of the diversity in a community given various PCR associated biases (Sharpton et al., 2011; Hong et al., 2009; Logares et al., 2013). Moreover, it can give considerably varying estimates of diversity due to the 16S gene variants of the bacterial genera targeted, the sequencing platform or the various centers of choice (A, B, C, D) inside each gene variant window, e.g V1-V3. (Jumpstart Consortium Human Microbiome Project Data Generation Working Group, 2012), whereas sequencing error and incorrectly assembled amplicons (i.e., chimeras) are often difficult to identify (Wylie et al., 2012) due to the presence of artifacts. Amplicon sequencing is limited to the analysis of taxa for which taxonomically informative genetic markers are known and can be amplified. Since the 16S locus can be transferred between distantly related taxa (i.e., horizontal gene transfer), analysis of 16S sequences for the estimation of the community diversity may result in overestimations (Acinas et al., 2004). Finally, the high level of its sequence conservation limits the power for resolving closely related organisms (Mende et al., 2013)

However, apart from 16S rRNA amplicon sequencing, it is actually possible to infer OTU’s based on raw metagenomic reads without PCR amplification, like in the case of the MGnify analysis pipeline version 4.1 (https://www.ebi.ac.uk/metagenomics/pipelines/4.1) (Mitchel et al. 2019). This version of the pipeline was used to analyse the datasets of *P. purpurea*, as they were deposited on 2017 at the Ebi metagenomics database. The process of grouping DNA sequences with at least 97% similarity is referred to as “OTU binning”. In this approach, the consensus sequence that is determined by the most common nucleotide at each position of the 16S ribosomal RNA gene provided in the raw data through the chosen sequencing platform, represents the different sequences that are assigned into OTU’s. The comparison of these OTU consensus sequences to reference databases of 16s rRNA, *e.g*. SILVA (http://www.arb-silva.de) or RDP (http://rdp.cme.msu.edu) provides the taxonomy assignment. This particular version of the pipeline (4.1) employs a novel tool (MAPseq) (Rodriguez et al., 2017) that was used in 2018 when the analysis of P*. purpurea* datasets deposited in MGnify was performed. This tool uses classified and unclassified reference sequences that are pre-clustered into OTU’s, which also hierarchically have different similarity cuttof’s so as to identify taxa of bacteria that are unknown. The hierarchical division of the OTU’s based on the sequence similarity of the 16S rRNA variants being assigned different cuttof values serves as an additional level of information that is used in tandem with the initial 97% cuttof to provide further resolution.

However, even metagenomic sequencing is not without challenges. In fact, metagenomic data is relatively complex and large, thus complicating the informatic analysis. For example, it can be difficult to determine the genome from which a read was derived, and most communities are so diverse that most genomes are not completely represented by reads. Furthermore, metagenomes tend to be relatively expensive to generate compared to amplicon sequences, especially in complex communities or when host DNA greatly outnumbers microbial DNA. However, ongoing advances in DNA sequencing technology are improving the affordability of metagenomic sequencing and significant steps have been made in that direction.

Here we report the metagenomic profiling of the microbial communities associated to *Porphyra purpurea* (Roth) C. Agardh, 1824 from four different localities. Sequences from eight metagenomic datasets of microbiota associated to the red algae *P. purpurea* have been quality controlled and pre-processed and then analyzed using MetaPhlAn 3.0. A qualitative and quantitative taxonomic profiling of the different bacterial communities at the species level has been performed.

A comparison of the bacterial diversity among the same locality and between the different ones is highlighted. Furthermore, the processed data from the publicly available datasets were juxtaposed with the results reported by MGnify database (www.ebi.ac.uk/metagenomics/).

Our analysis will provide further insight to the scantly known bacterial communities associated with the species *P. purpurea*, using a method for shotgun metagenomic data analysis that could be also applied downstream to the investigation of the functional profiling of the most abundant metabolic pathways of the host-associated bacterial communities identified.

## 2. Materials and methods

### 2.1 Datasets

The datasets analysed in this study originated from algal blades of *P. purpurea* collected from four different localities: United Kingdom, Ireland, USA and Italy. Specimens of *P. purpurea* from United Kingdom were collected at Sidmouth on January 2015, (50.677 N; 3.240 W). Ireland sampling was carried out within Bantry bay (51.640 N; 9.710 W) on November 2014. *P. purpurea* was collected in USA along the atlantic shores of Maine (44.342 N; 68.065 W) on January 2015, while the sampling in Italy was carried out along the Northwestern coasts of Sardinia (41.162 N; 8.622 E) on February 2015.

All samples were sequenced at DOE Joint Genome Institute on Illumina HiSeq 2500 platform following the shotgun metagenomic illumina protocol using forward and reverse TruSeq adapters resulting in paired end reads. The sequences were obtained from EMBL-EBI European Nucleotide Archive database on 10th May 2020. For Maine (USA), two datasets were available, with 24.8 Gb and 80 million reads for Maine_Asex_4 and 16.2 Gb and 53.7 million reads for Maine_Asex2. For Italy two datasets were available, Italy_Porto_4 with 27.9 Gb and 92 million reads, Italy_Porto_5 20.5 Gb and 67 million reads. The Irish dataset had 28.5 Gb and 94 million reads. Concerning the three datasets from United Kingdom, Sidmouth_Asex_1 had 16.5 Gb and 54.5 million reads, Sidmouth_Asex_2 24 Gb and 79.5 million reads and Sidmouth_Male_2 18.9 Gb and 62.6 million reads.

### 2.2 Pre-processing

The reads were assessed using FastQC (version 0.11.8) (Andrews, 2010). The assessment evaluated the overall quality of the sequenced reads, their average length, the duplicates found in each sample as well as provided an overview of the GC content present in each sample. The quality and the other metrics were used as a guide for the required parameterization of the metagenomic profiling procedure. Trimmomatic (version 0.39) (Bolger et al., 2014) was used to remove the TruSeq illumina adapters after the addition of their sequence in a custom database.

### 2.3 Metagenomic profiling

The pre-processed reads were uploaded to HPC server “Zorbas” of HCMR (Hellenic Center of Marine Research) located at Heraklion (Crete), where the analyses took place. MetaPhlAn 3.0 (Truong et al., 2015) was used for the metagenomic profiling of the reads using a stat q value of 0.1 along with the default configurations. For each dataset the configuration used was exactly the same. It must be noted that one dataset from the locality of Sidmouth (“Sidmouth Male 1”) did not provide any results whatsoever and was therefore excluded from the analysis. The ChocoPhlan pan-genome database v30 was used for the purpose of the taxonomy identification to species level with MetaPhlAn 3.0 (mpa_v30_CHOCOPhlAn_201901_marker_info.txt.bz2) (profiled metagenome bowtie2 output in Supplementary File 1).

### 2.4 Post-profiling analysis

Clade specific markers, calculation of the number of aligned reads and all the markers found in the metagenomic profiles were extracted from the primary output files provided by MetaPhlAn 3.0 (see Supplementary File 1). The clade specific markers as well as the marker count (for complete taxonomies to species level exclusively) of the metagenomic profiling were used to compare the results presented here with the results of the MGnify database (Mitchell et al., 2020) where the same datasets are deposited and analysed via the method of 16s ribosomal RNA gene (Table 1).

**Table 1.**
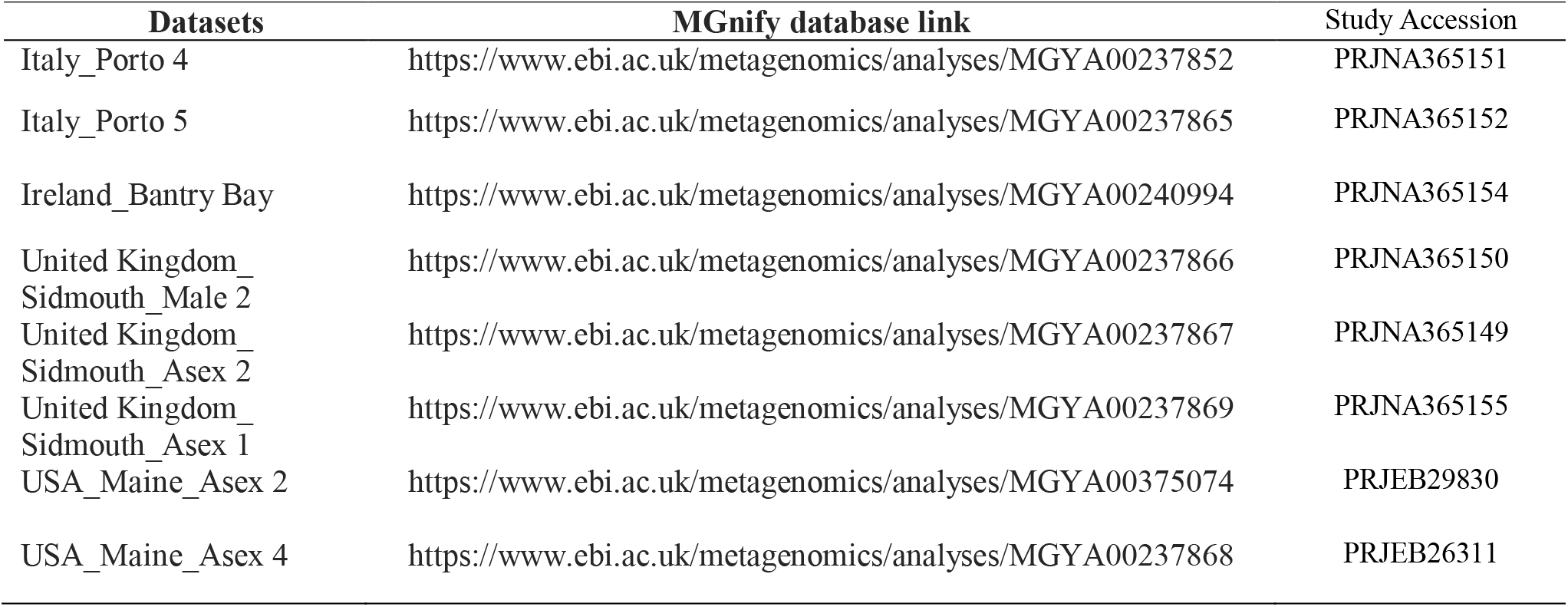
MGnify database links for each analysed dataset.

| Datasets | MGnify database link | Study Accession |
| --- | --- | --- |
| Italy_Porto 4 | https://www.ebi.ac.uk/metagenomics/analyses/MGYA00237852 | PRJNA365151 |
| Italy_Porto 5 | https://www.ebi.ac.uk/metagenomics/analyses/MGYA00237865 | PRJNA365152 |
| Ireland_Bantry Bay | https://www.ebi.ac.uk/metagenomics/analyses/MGYA00240994 | PRJNA365154 |
| United Kingdom_<br>Sidmouth_Male 2 | https://www.ebi.ac.uk/metagenomics/analyses/MGYA00237866 | PRJNA365150 |
| United Kingdom_<br>Sidmouth_Asex 2 | https://www.ebi.ac.uk/metagenomics/analyses/MGYA00237867 | PRJNA365149 |
| United Kingdom_<br>Sidmouth_Asex 1 | https://www.ebi.ac.uk/metagenomics/analyses/MGYA00237869 | PRJNA365155 |
| USA_Maine_Asex 2 | https://www.ebi.ac.uk/metagenomics/analyses/MGYA00375074 | PRJEB29830 |
| USA_Maine_Asex 4 | https://www.ebi.ac.uk/metagenomics/analyses/MGYA00237868 | PRJEB26311 |

### 2.5 Visualization

Hclust2 was used to produce the heatmaps, while GraPhlAn (Asnicar et al., 2015) was applied for the graphical representation of the bacterial taxonomies found according to their relative abundances.

## 3. Results and Discussion

The bacterial species identified from the eight datasets are depicted in Fig. 1, according to their relative abundances. A total of 43 bacterial species were identified, with the more prominent ones being *Psychrobacter JCM 18900, Pseudoalteromonas distincta, Pseudoalteromonas arctica, Pseudoalteromonas BSi20652, Staphylococcus pasteuri, Staphylococcus epidermidis, Staphylococcus warneri, Bacillus pumilus, Bacillus altitudinis, Psychrobacter P11G3, Dokdonia donghaensis, Cellulophaga baltica, Bacillus stratosphericus. Cutibacterium acnes* and *Pseudoalteromonas distincta* (Fig. 1, Table 1)

**Figure 1.**
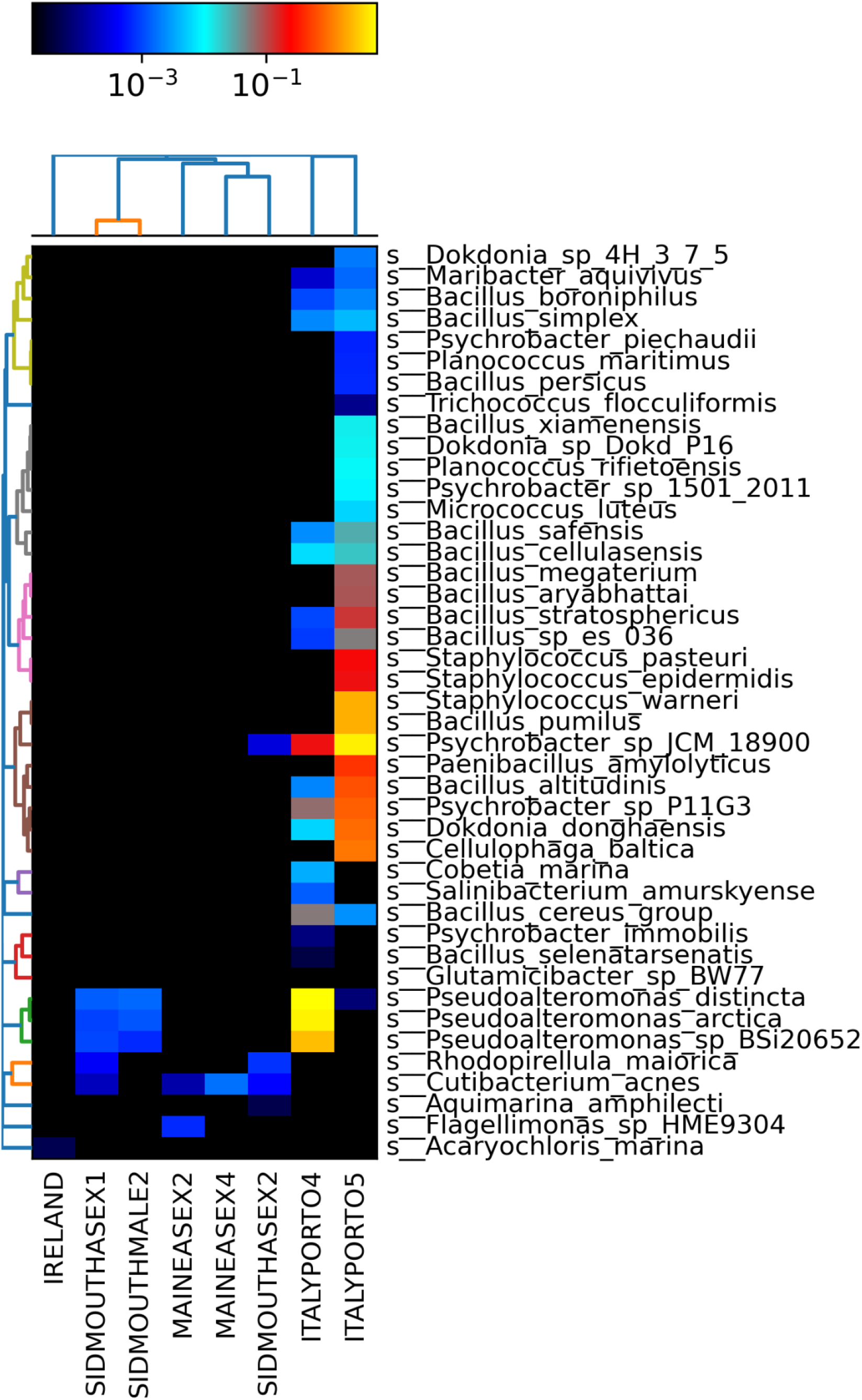
Heatmap of the identified bacterial species according to their respective datasets based on their relative abundance values as calculated from MetaPhlAn 3.0.

In detail, *P. distincta* was found in four datasets (Sidmouth_Male 2, Sidmouth_Asex 1, Italy_Porto 4, and Italy_Porto 5), and *C. acnes* was found in four datasets as well (Sidmouth_Asex 1, Sidmouth_Asex 4, Maine Asex_2, Maine Asex 2). *B. altitudinis, P*. P11G3 *D*. d*onghaensis* and *B*. s*tratosphericus* were found in two datasets (Italy_Porto 4 and Italy Porto_5), whereas *S. pasteuri, S. warneri, S. epidermidis B. pumilus*, and *C. baltica* were found only in one dataset (Italy Porto 5). *Psychrobacter JCM 18900* was found in three datasets (Sidmouth_Asex 1, Italy_Porto 4, and Italy_Porto 5), such as *P. arctica* and *P. BSi20652* (Sidmouth_Asex 1, Sidmouth_Male 2, Italy_Porto 4). *Glutamibacter BW77* was found in the lowest relative abundance with respect to the other species in Italy_Porto 4 dataset.

The bacterial diversity associated to *P. purpurea* datasets from Italy was spread across five phyla on the total of six phyla identified, with a total of 38 species (Table 2). The dataset from Ireland showed just one species not found in any other area, the cyanophean *Acaryochloris marina*. The datasets from Maine included two species *Cutibacterium acnes* (found also in the dataset from Italy and UK) and *Flagellimonas* sp. HME9304. Finally, the datasets from Sidmouth (United Kingdom) included seven species among three classes, Actinobacteria, Flavobacteria, Planctomycetia and Gamma-proteobacteria.

**Table 2.**
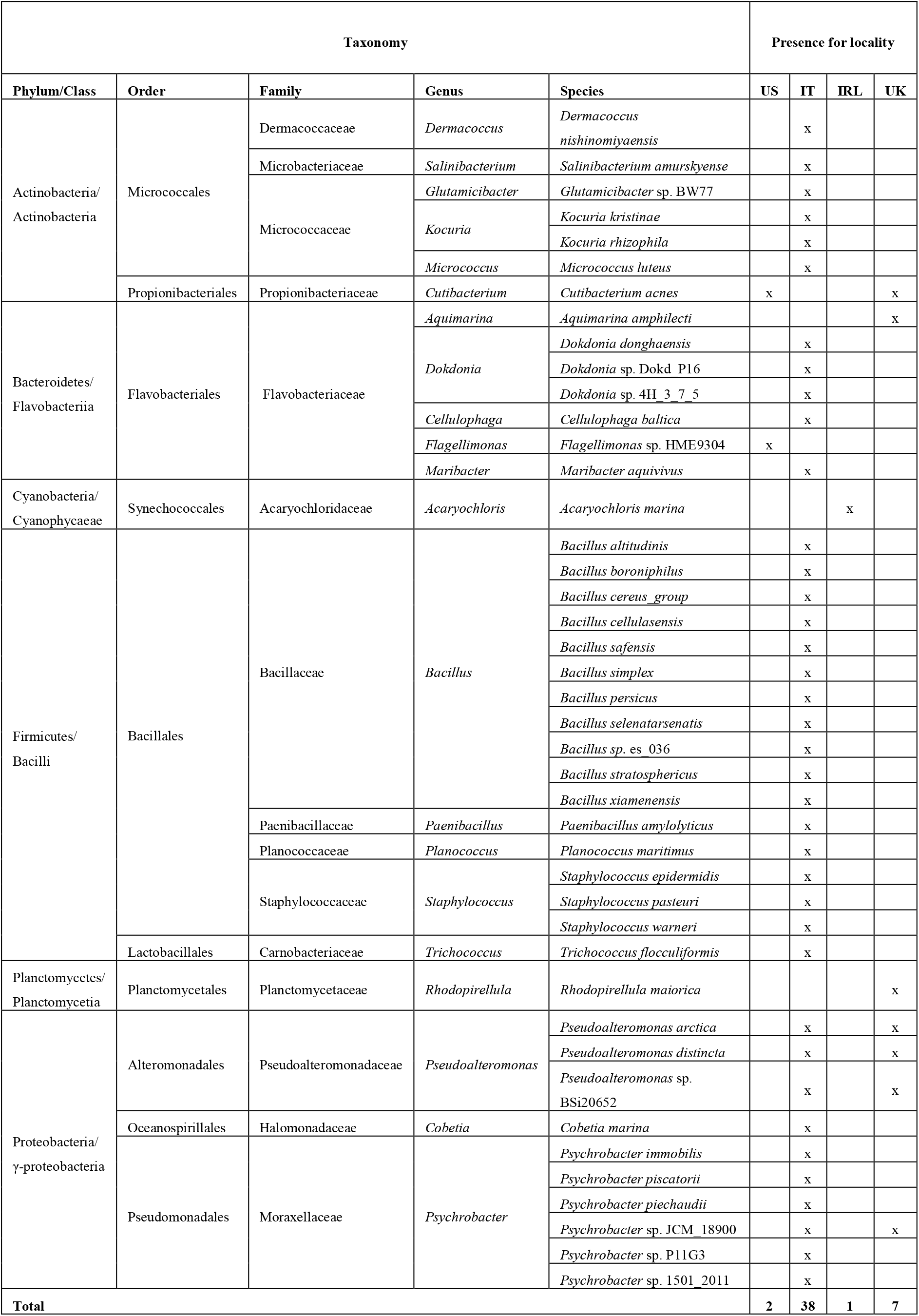
List of the bacterial taxonomical diversity in the four localities (US, USA; IT, Italy; IRL, Ireland; UK, United Kingdom).

The complete bacterial taxonomical diversity found in the metagenomic profiling analysis based on their relative abundance values is shown in Fig. 2. The genera *Pseudoalteromonas, Psychrobacter, Bacillus* and *Staphylococcus* are the most prominent ones with *Dokdonia* and *Cellulophaga* being the less prominent of the most abundant genera identified. The relative abundances estimated are normalized according to the each dataset’s read count, so as to be merged together and presented as a whole in graphlan’s circular representation in Fig. 2. The depiction of Fig. 2 results are furthermore elucidated and verified in the section 3.1 where the putative taxa characterised have been compared with existing literature. The genus *Staphylococcus* represent an exception and further investigation are needed to explain the presence of this pathogenic group of bacteria associated with macro-algae.

**Figure 2.**
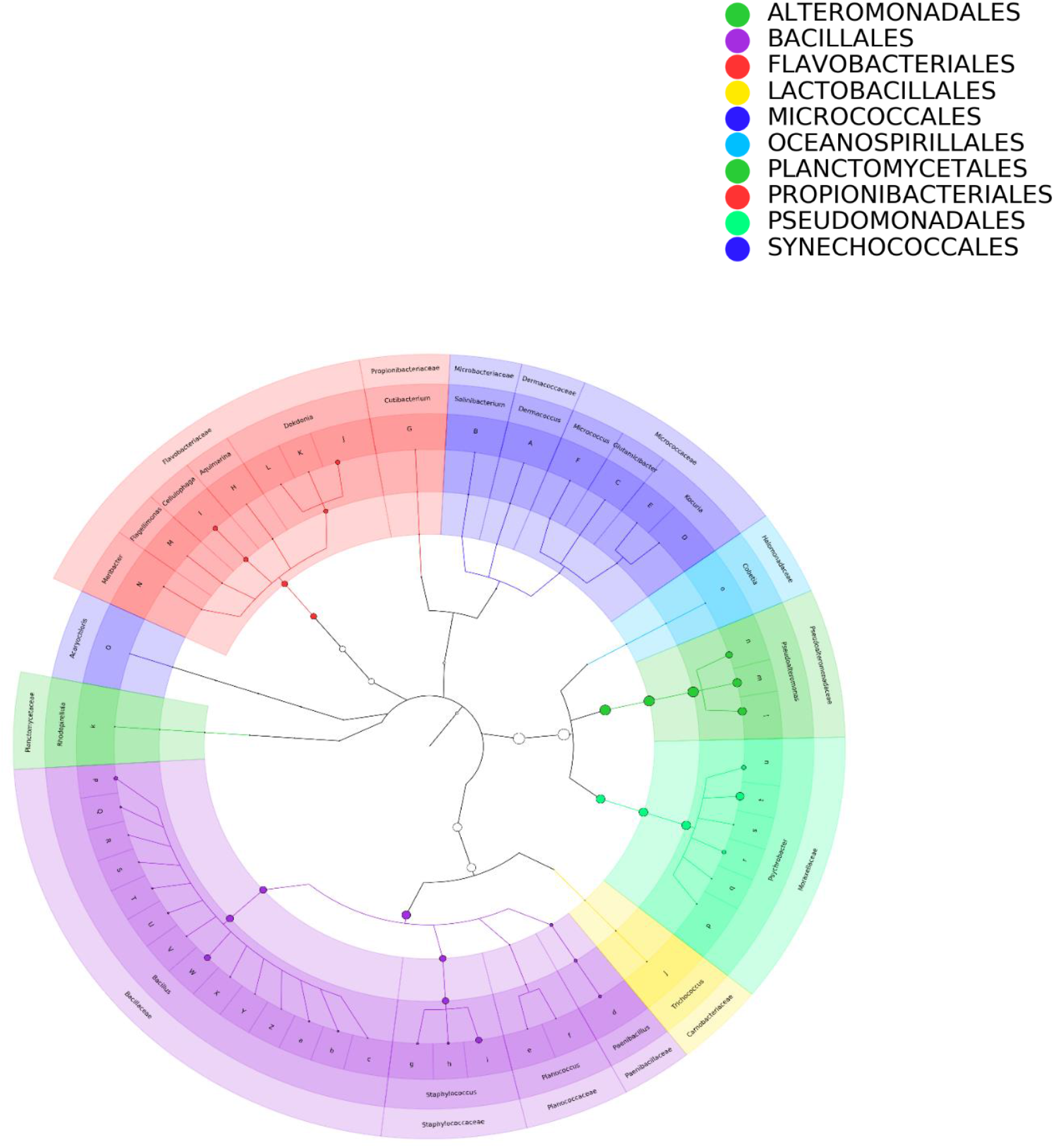
Circular representation of identified bacterial taxonomies according to their relative abundances. The nodes of the clades correspond to the different taxonomic levels with the lowercase letters indicating the different species.

The common species identified by both methods were five as shown in the heatmap shown in Fig. 3, *Acaryochloris marina, Dokdonia donghaensis, Planococcus maritimus, Psychrobacter P11G3, Staphylococcus epidermidis*. Additionally, at higher taxonomic level the two methods shared the genera *Glutamicibacter, Bacillus, Pseudoalteromonas, Psychrobacter, Cellulophaga, Dokdonia, Maribacter, Paenibacillus* and *Rhodopirellula* (out of 21 found with MetaPhlAn 3.0). Since MetaPhlAN 3.0 provides relative abundances in its default configuration, it is not possible to compare with results provided by the MGnify pipeline directly. For this reason, the counts of the mapped reads to the reference pan-genome database used with MetaPhlAn were extracted and summed. These counts can be referred to as pan-genome counts, or “pan-genome based OTU counts” and since they are clade specific, can be compared to OTU counts. In Fig. 3 purposes all SSU have been summed (coding area of small ribosomal DNA gene) and LSU (long subunit) counts for the species that were commonly identified by both of them and kept the uniquely identified species counts from either SSU or LSU separate.

**Figure 3.**
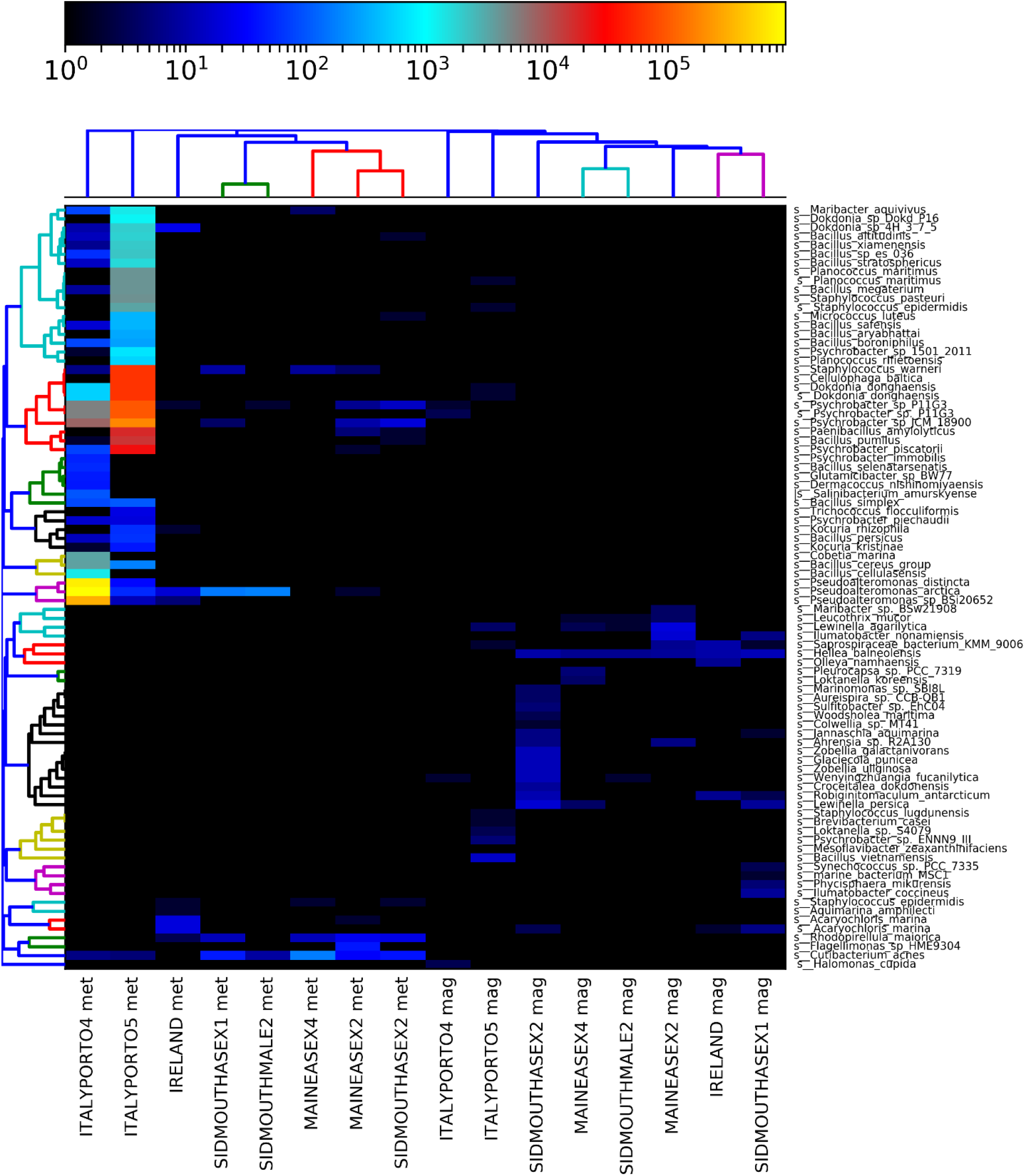
Comparison at species level of the MetaPhlAn 3.0 (indicated by “met”) to the 16S rRNA results from the MGnify database (indicated by “mag”). The comparison was made with the OTU counts for the identified bacterial species and the “OTU” analogous counts from the metagenomic profiling procedure using MetaPhlAn 3.0. The counts from MetaPhlAn were extracted from the output bowtie2 files and include only the counts from the species that are unequivocally detected. These species are the ones that had reads mapped to the reference database for all the taxonomical levels.

### 3.1 Comparisons with previous studies

Literature regarding host-associated macro algal bacteria generally provide information up to phylum level (Singh and Reddy, 2013), albeit the most reported orders are also found in this study, such as Actinobacteria, Bacteroidetes (Aires et al., 2013; Tujula et al., 2010), Flavobacteria (Matsuo et al., 2005), Gammaproteobacteria and Planctomycetes (Burke et al. 2011; Bondoso et al., 2013; Meusnier et al., 2001). The datasets analysed report Proteobacteria in United Kingdom and Italy, Actinobacteria in USA, United Kingdom and Italy, Bacteroidetes in Italy and USA, Firmicutes in Italy, Planctomycetes in United Kingdom and Cyanobacteria in Ireland.

The genus *Pseudoalteromonas* resulted abundant in three datasets. This could be explained by the fact that heterotrophic marine bacteria have developed abilities to grow on the macroalgal polysaccharides that constitute half of seaweed biomass (Gobet et al., 2018). Indeed, the species *Pseudoalteromonas carrageenovora* 9^T^ (ATCC 43555^T^), a marine gammaproteobacterium, has a diverse repertoire of carbohydrate-active enzymes (CAZymes), mainly for the degradation of macroalgal polysaccharides (laminarin, alginate, FCSP, carrageenans) (Gobet et al., 2018).

The genus *Bacillus* was found in the Italian datasets in considerabe relative abundances. Many of the studies done on epiphytic bacteria of macroalgae are oriented towards evaluating antibacterial activity, which is more than justified if we consider that the surface of the macroalgae is an important source of substrate and nutrients, leading in a competitive environment (Armstrong et al., 2001). *Bacillus* and *Pseudoalteromonas* genera have been correlated with antimicrobial activity in contrast to four other macroalgal associated strains *Bacillus algicola*, *Formosa algae* and *Algicola bacteriolytica* (Goecke et al., 2013). In the same study three *Bacillus* species were identified from brown and red macroalgal species, that are common with our findings, *Bacillus* s*afensis, B. altitudinis, B. cereus*. *Bacillus algicula* was also isolated from thallus of *Fuscus evanescens* (Ivanova et al., 2004).

In our datasets the genus *Psychrobacter* resulted highly represented. Indeed, psychrotrophic bacteria typically grow at low temperatures and the samplings were all carried out during winter months. Similar bacterial strains (16S rRNA), such as *Psycrobacter aquimaris, P. celeris, P. nivimaris, Psychromonas arctica*, were also isolated from brown algae *Undaria pinnatifida* (Lee et al., 2006), whose sporophytes are cultured during winter.

Furthermore, *Alphaproteobacteria* and *Gammaproteobacteria* were also isolated from the Australian red algae *Amphiroa anceps*, while *Bacteroidetes* and *Gammaproteobacteria* were isolated from another red algae, *Corallina officinalis* (Brodie et al., 2016). Burke et al. (2011) employed metagenomic analysis of red alga *U. australis* associated bacterial communities and found that it predominantly consisted of sequences from *Proteobacteria* (64.0%), *Bacteroidetes* (27.6%) and *Planctomycetes* (3.4%).

With regards to the results obtained from *P. purpurea* datasets, bacterial communities sampled in the different localities were not comparable, with the Italian datasets characterized by a richer biodiversity and the Irish one the lowest. Despite the fact that macroalgae belonging to the same species occurring in different geographical locations generally have similar bacterial communities to those from different species in the same ecological niche (Lachnit et al., 2009), some studies reported that the composition of bacterial phyla can vary among replicates of the same algal species and also within the same season (Lachnit et al., 2011).

### 3.2 Methodological insights

Metagenomic analyses can be challenging, mainly due to the volume and nature of the sequencing data. Whole genome shotgun sequencing (WGS) runs can yield hundreds of millions of sequences and these sequences are usually relatively short ranging from approximately 100 to more than 300 base pairs for typical Illumina paired-end runs, which is the dominant sequencing platform for metagenomics. The large size of metagenomic datasets as well as their nature require effective approaches that can provide unambiguous results. Our method of choice infers the presence and read coverage of clade specific markers to unequivocally detect the taxonomic clades present in a microbiome sample and estimate their relative abundance. MetaPhlAn 3.0 includes an expanded set of markers for each bacterial species originating from ~99,500 bacterial genomes following the pan-genome rationale. Pan, from the Greek word (*παν*, meaning whole) (Tettelin et al., 2005), which includes a core genome comprised of genes present in all strains, a “redundant” genome containing genes not present in one or more strains and genes that are unique to each strain. Studies of pan-genomes arose from these new possibilities and reflect the notion of bacterial species more accurately (Medini et al., 2005; Ramasamy et al., 2014) in this way the limits and intraspecific variation among bacteria, and microbial species in general, are drawn from the unique repertoire of genes of a group of organisms thus making the taxonomical classification more precise and reliable. MetaPhlAn characterizes metagenomic marker gene homologs by similarity binning, which classifies a read into a taxonomic or phylogenetic group based on its similarity to previously identified genes or proteins. It uses an extensive database of phylogenetic clade-specific markers (i.e., families that are single copy and generally only common to a monophyletic group of taxa) to assign metagenomic sequences to specific taxonomic groups (Segata et al., 2012). This extensive database and its marker specificity, with hundreds of unique markers for each species, provides the core potential of accurate and less ambiguous identification. Furthermore, MetaPhlAn 3.0 has increased functionality compared to the previous version, such as internal scoring methods for each step, so as to evaluate the intermediate data being produced until the end results. The inherent mapq function plays that role, which discards mapped reads that have a value less than 5. Moreover, due to the minimum read length option, all reads with 70 or less bases are discarded as well, according to the default configuration. For each clade, MetaPhlAn species relative abundances are calculated by creating a vector of markers and their count (number of reads hitting the marker). According to the quantile value chosen, e.g stat_q value of 0.1, the markers taken into consideration for the local abundance calculation are the ones falling between the 10th and 90th percentile and using this subset of markers, the local abundance is calculated by dividing the sum of the number of reads mapping to the clade markers by the sum of the clade markers length. The number of reads mapping to the clade is estimated by multiplying the local clade abundance by the clade’s average genome size. Then, relative abundances are calculated as local clade abundance divided by the sum of all the local abundances. The normalization procedure is where the genome size variability among different bacterial strains or species is taken into consideration, and the relative abundance reflects more accurately the multitude and length of the markers that the reads are mapped to, so as to account for their natural variability.

A main approach in metagenomic analyses is the assembly of the sequenced data into a metagenome. The reason why contigs are not made from the reads in MetaPhlAn (there is not an assembled metagenome) is because in WGS sequencing the reads are often <150 nt and the fragments being sequenced often >300 nt. So, the reads, often, do not overlap enough to be assembled into contigs. Whereas in 16S sequencing the 16S gene amplification primers and the method of sequencing are designed to produce overlapping paired-end reads in order to be able to assemble them. In MetaPhlAn the reads are treated as independent single reads for mapping to the reference database. Paired-end reads are expected to align to the reference at more-or-less a fixed distance apart and in opposite orientation. These ‘expectations’ are ignored.

It is of importance to note that all of the datasets used here had reads with average length approximating to 150 nt as well as high phred quality scores. High quality datasets increase the probability of a less biased and more reliable study. The objectives of this study were to accurately profile the putative host associated bacterial species *P. purpurea*, to evaluate the results comparing them to existing literature of bacteria species associated with macro algae and to give insights concerning the methodologies adopted. In this study, the 16S rRNA based OTUs provide different results by using the 4.1 version of MGnify pipeline. This could be explained by the fact that it exploits just a gene that has a slow evolutionary rate and shows some polymorphism, having nine variants (V1-V9) in bacterial species, leading in low resolution between closely related taxa. On the other hand, MetaPhlAn 3.0 increased its reference database from ~13.500 bacterial genomes to ~99.500 and the marker genes to ~1.1 million thus increasing its resolving capacity and the number of identifiable taxa considerably. Additionally, the option of estimation of the unknown portion of the dataset was added making the relative abundance calculation more precise. Thus, we propose the pan-genome approach and MetaPhlAn as an analytical tool, which shows good resolution for the studies of bacterial communities.

### 3.1 Conclusion

- 43 bacterial species were identified in eight datasets from four localities.
- The pan-genome approach used in this study can effectively resolve taxonomical identification of macroalgae bacteria-associated species. MetaPhlAn 3.0 showed high efficiency and resolution.
- The expected difference in taxonomical resolution between a16S rRNA OTU based method for analysing raw reads, as well as the contradicting results, verifies the agreement with other studies (Truong et al. 2015). This was made more evident due to the fact that the two approaches shared nine out of the total 21 genera identified with MetaPhlAn 3.0, but only five of 43 species.
- The phyla of the profiled bacterial species are found in the majority of the published literature of epiphytic/endophytic bacteria of macroalgal species.
- A variability in composition among the datasets and localities has been identified. Potential alternative causes might be: sampling contamination, environmental conditions, sequencing contamination, taxon sampling issues.

## References

Acinas, S.G., Marcelino, L.A., Klepac-Ceraj, V., Polz, M.F., 2004. Divergence and redundancy of 16S rRNA sequences in genomes with multiple Rrn operons. J. Bacteriol. 186, 2629–2635.

Andrews, S., 2010. FastQC: a quality control tool for high throughput sequence data.

Aires, T., Serrao, E.A., Kendrick, G., Duarte, C.M., Arnaud-Haond, S., 2013. Invasion is a community affair: clandestine followers in the bacterial community associated to green algae, *Caulerpa racemosa*, track the invasion source. PLoS One 8 e68429.

Asnicar, F., Weingart, G., Tickle, T.L., Huttenhower, C., Segata, N., 2015. Compact graphical representation of phylogenetic data and metadata with GraPhlAn. PeerJ 3, e1029.

Armstrong, E., Yan, L., Boyd, K.G., Wright, P.C., Burgess, J.G., 2001. The symbiotic role of marine microbes on living surfaces. Hydrobiologia 461, 37–40.

Bolger, A. M., Lohse, M., Usadel, B., 2014. Trimmomatic: a flexible trimmer for Illumina sequence data. Bioinformatics 30, 2114–2120.

Bondoso, J., Balague, V., Gasol J.M., Lage O.M., 2013. Community composition of the Planctomycetes associated with different macroalgae. FEMS microbiology ecology, 88, 445–456.

Biller, S.J., Berube, P.M., Dooley, K., Williams, M., Satinsky, B.M., Hackl, T., Hogle, S.L., Coe, A., Bergauer, K., Bouman H.A., Browning, T.J, De Corte, D., Hassler, C., Hulston, D., Jacquot, J.E., Maas, E.W., Reinthaler, T., Sintes, E., Yokokawa, T., Chisholm, S.W., 2018. Marine microbial metagenomes sampled across space and time. Sci. Data 5, 180176.

Bowen De León, K., Gerlach, R., Peyton, B.M., Fields, M.W., 2013. Archaeal and bacterial communities in three alkaline hot springs in Heart Lake Geyser Basin, Yellowstone National Park. Front. Microbiol. 4, 330.

Brodie, J. Irvine, L.M., 2003. Seaweeds of the British Isles. Vol. 1 Part 3B. Bangiophycidae. Intercept, Hampshire.

Brodie, J., Williamson, C., Barker, G. L., Walker, R. H., Briscoe, A., Yallop, M., 2016. Characterising the microbiome of *Corallina officinalis*, a dominant calcified intertidal red alga. FEMS microbiology ecology 92, fiw110.

Burke, C., Steinberg, P., Rusch, D., Kjelleberg, S., Thomas, T., 2011. Bacterial community assembly based on functional genes rather than species. P. Natl. Acad. Sci. USA 108, 14288–14293.

Gantt, E., Mine Berg, G., Bhattacharya, D., Blouin, N.A., Brodie J.A., Xin Chan C., Collén, J., Cunningham Jr, F.X., Gross, J., Grossman, A.R., Karpowicz, S., Kitade, Y., Klein, A.S., Levine, I.A., Lin, S., Lu, S., Lynch, M., Minocha, S.C., Müller, K., Neefus, C.D., Cabral de Oliveira, M., Rymarquis, L., Smith, A., Stiller, J.W., Wu W., Yarish, C., Zhuang, Y., Brawley, S.H., 2010. *Porphyra:* complex life histories in a harsh environment: *P. umbilicalis*, an intertidal red alga for genomic analysis. In: Seckbach J., Chapman D. (eds) red algae in the genomic age. Cellular Origin, Life in Extreme Habitats and Astrobiology, vol 13. Springer, Dordrecht.

Gobet, A., Barbeyron, T., Matard-Mann, M., Magdelenat, G., Vallenet, D., Duchaud, E., Michel, G., 2018. Evolutionary evidence of algal polysaccharide degradation acquisition by *Pseudoalteromonas carrageenovora* 9T to adapt to macroalgal niches. Frontiers in Microbiology 9, 2740.

Goecke, F., Thiel, V., Wiese, J., Labes, A., Imhoff, J.F., 2013. Algae as an important environment for bacteria-phylogenetic relationships among new bacterial species isolated from algae. Phycologia 52, 14–24.

Hong, S., Bunge, J., Leslin, C., Jeon, S., Epstein, S.S., 2009. Polymerase chain reaction primers miss half of rRNA microbial diversity. ISME J. 3, 1365–1373.

Hugenholtz, P., Pace, N. R., 1996. Identifying microbial diversity in the natural environment: a molecular phylogenetic approach. Trends Biotechnol. 14, 190–197.

Ivanova, E.P., Alexeeva, Y.A., Zhukova, N.V., Gorshkov, A N.M., Buljan, V., Nicolau, D.V., Mikhailov, V.V., Christen, R., 2004. *Bacillus algicola* sp. nov., a novel filamentous organism isolated from brown alga *Fucus evanescens*. Systematic and applied microbiology 27, 301–307.

Jumpstart Consortium Human Microbiome Project Data Generation Working Group, 2012. Evaluation of 16S rDNA-based community profiling for human microbiome research. PLoS ONE 7, e39315.

Konopka, A., 2009. What is microbial community ecology? The ISME journal, 3, 1223–1230.

Lachnit, T., Blümel, M., Imhoff, J. F., Wahl, M., 2009. Specific epibacterial communities on macroalgae: phylogeny matters more than habitat. Aquatic Biology 5, 181–186.

Lachnit, T., Meske, D., Wahl, M., Harder, T., Schmitz, R., 2011. Epibacterial community patterns on marine macroalgae are host-specific but temporally variable. Environmental microbiology 13, 655–665.

Lee, Y.K., J.J. Hyun, Hong K.L., 2006. Marine bacteria associated with the Korean brown alga, *Undaria pinnatifida*. Korean J. Microbiol. 44, 694–698.

Lozupone, C.A., Knight, R., 2007. Global patterns in bacterial diversity. Proc. Natl. Acad. Sci. U.S.A. 104, 11436–11440.

Logares, R., Sunagawa, S., Salazar, G., Cornejo-Castillo, F.M., Ferrera, I., Sarmento, H., Hingamp, P., Ogata, H., de Vargas C., Lima-Mendez, G., Raes, J., Poulain, J., Olivier Jaillon, O., Patrick Wincker, P., Kandels-Lewis, S., Karsenti, E., Bork, P., Acinas, S.G., 2013. Metagenomic 16S rDNA illumina tags are a powerful alternative to amplicon sequencing to explore diversity and structure of microbial communities. Environ. Microbiol. 16, 2659–2671.

Matsuo, Y., Imagawa, H., Nishizawa, M., Shizuri, Y., 2005. Isolation of an algal morphogenesis inducer from a marine bacterium. Science 307, 1598.

McCliment, E.A., Voglesonger, K.M., O’Day, P.A., Dunn, E.E., Holloway, J.R. Cary, S.C., 2006. Colonization of nascent, deep-sea hydrothermal vents by a novel Archaeal and Nanoarchaeal assemblage. Environ. Microbiol. 8, 114–125.

Medini, D., Donati, C., Tettelin, H., Masignani, V., Rappuoli, R., 2005. The microbial pangenome. Curr Opin Genet Dev 15, 589–94.

Mende, D., Sunagawa, S., Zeller, G. Bork, P., 2013. Accurate and universal delineation of prokaryotic species. Nat. Methods 10, 881–884. https://doi.org/10.1038/nmeth.2575

Meusnier, I., Olsen, J.L., Stam, W.T., Destombe, C., Valero, M., 2001. Phylogenetic analyses of *Caulerpa taxifolia* (Chlorophyta) and of its associated bacterial microflora provide clues to the origin of the Mediterranean introduction. Mol. Ecol. 10, 931–946.

Milanese, A., Mende, D.R., Paoli, L., Salazar, G., Ruscheweyh, H., Cuenca, M., Hingamp, P., Renato Alves, R., Costea, P.I., Coelho L., Schmidt, T.S.B., Almeida A., Mitchell, A.L., Finn, R.D., Huerta-Cepas, J., Bork, P., Zeller, G., Sunagawa, S., 2019. Microbial abundance, activity and population genomic profiling with mOTUs2. Nat. Commun. 10, 1014.

Mitchell, A.L., Almeida, A., Beracochea, M., Boland, M., Burgin, J., Cochrane, G., Crusoe, M.R., Kale, V., Potter, S.C., Richardson, L.J., Sakharova, E., Scheremetjew, M., Korobeynikov, A., Shlemov, A., Kunyavskaya, O., Lapidus, A., Finn, R.D., 2020. MGnify: the microbiome analysis resource in 2020. Nucleic Acids Research 48, 570–578.

Mori, S., Yamazaki, A., Matsuyama-Serisawa, K., Fukuda, S., Mizuta, H., Saga, N., 2004. Effect of two symbiotic bacteria for growth of *Porphyra yezoensis* (Rhodophyta, Bangiales) in axenic culture. Aquaculture Science 52, 239–244.

Namba, A., Shigenobu, Y., Kobayashi, M., Kobayashi, T., Oohara, I., 2010. A new primer for 16S rDNA analysis of microbial communities associated with *Porphyra yezoensis*. Fisheries Science 76, 873–878.

Pace, N. R., 1997. A molecular view of microbial diversity and the biosphere. Science 276, 734–740.

Pace, N.R., Stahl, D.A., Lane, D.J., Olsen, G.J., 1986. The analysis of natural microbial populations by ribosomal RNA sequences. In Advances in microbial ecology. Springer, Boston, MA.

Ramasamy, D., Mishra, A.K., Lagier, J.C., Padhmanabhan, R., Rossi M., Sentausa E., Raoult, D., Fournier P., 2014. A polyphasic strategy incorporating genomic data for the taxonomic description of novel bacterial species. Int. J. Syst. Evol. Microbiol. 64, 384–91.

Rappé, M.S., Giovannoni, S.J., 2003. The uncultured microbial majority. Annu. Rev. Microbiol. 57, 369–394.

Matias Rodrigues, J.F., Schmidt, T.S.B., Tackmann, J., von Mering, C., 2017. MAPseq: highly efficient k-mer search with confidence estimates, for rRNA sequence analysis. Bioinformatics 33, 3808–3810. doi:10.1093/bioinformatics/btx517

Saga, N., Kitade, Y., 2002. *Porphyra*: a model plant in marine sciences. Fisheries science, 68, 1075–1078.

Segata, N., Waldron, L., Ballarini, A., Narasimhan, V., Jousson, O., Huttenhower, C., 2012. Metagenomic microbial community profiling using unique clade-specific marker genes. Nat. Methods 9, 811–814.

Sharpton, T.J., Riesenfeld, S.J., Kembel, S.W., Ladau, J., O’Dwyer, J.P., Green, J.L., 2011. PhylOTU: a high-throughput procedure quantifies microbial community diversity and resolves novel taxa from metagenomic data. PLoS Comput. Biol. 7, e1001061.

Singh, R.P., Reddy, C.R.K., 2014. Seaweed–microbial interactions: key functions of seaweed-associated bacteria. FEMS microbiology ecology 88, 213–230.

Soo, R.M., Wood, S.A., Grzymski, J.J., McDonald, I.R., Cary, S.C., 2009. Microbial biodiversity of thermophilic communities in hot mineral soils of Tramway Ridge, Mount Erebus, Antarctica. Environ. Microbiol. 11, 715–728.

Tang, Y.Z., Kang, Y., Berry, D., Gobler, C.J., 2014. The ability of the red macroalga, *Porphyra purpurea* (Rhodophyceae) to inhibit the proliferation of seven common harmful microalgae. Journal of applied phycology 27, 531–544.

Tettelin, H., Masignani, V., Cieslewicz, M.J., Donati, C., Medini, D., Ward N.L., Angiuoli, S.V., Crabtree, J., Jones A.L., Scott Durkin A., DeBoy, R.T., Davidsen, T.M., Mora, M., Scarselli, M., Margarit y Ros, I., Peterson, J.D., Hauser, C.R., Sundaram, J.P, Nelson, N.C., Madupu, R., Brinkac, L.M., Dodson, R.J., Rosovitz, M.J., Sullivan, S.A., Daugherty, S.C., Haft, D.H., Selengut, J., Gwinn, M.L., Zhou, L., Zafar, N., Khouri, H., Radune, D., Dimitrov, G., Watkins, K., O’Connor, K.J.B., Smith, S., Utterback, T.R., White, O., Rubens, C.E., Grandi, G., Madoff, L.C., Kasper, D.L., Telford J.L., Wessels, M.R., Rappuoli, R., Fraser, C.M., 2005. Genome analysis of multiple pathogenic isolates of *Streptococcus agalactiae*: implications for the microbial“pan-genome”. Proc. Natl. Acad. Sci. USA 102, 13950–13955.

Truong, D., Franzosa, E., Tickle, T.M., Weingart, G., Pasolli, E., Tett, A., Huttenhower, C., Segata, N., 2015. MetaPhlAn2 for enhanced metagenomic taxonomic profiling. Nat. Methods 12, 902–903.

Tujula, N.A., Crocetti, G.R., Burke, C., Thomas, T., Holmstrom, C., Kjelleberg, S., 2010. Variability and abundance of the epiphytic bacterial community associated with a green marine Ulvacean alga. ISME J4 301–311.

Wylie, K.M., Truty, R.M., Sharpton, T.J., Mihindukulasuriya, K.A., Zhou, Y., Gao, H., Sodergren, E., Weinstock, G.M., Pollard, K.S., 2012. Novel bacterial taxa in the human microbiome. PLoS ONE 7, e35294.

Yarish, C., Pereira, R., 2008. Mass production of marine macroalgae, In S.E. Jørgensen and B.D. Fath (eds.) Ecological Engineering. Vol. 3. Encyclopedia of Ecology. Elsevier, Oxford 2236–2247.

